# Comparative genomics and integrated system biology approach unveiled undirected phylogeny patterns, mutational hot spots, functional patterns and molecule repurposing for monkey pox virus

**DOI:** 10.1101/2022.12.21.521441

**Authors:** Nirjara Singhvi, Chandni Talwar, Utkarsha Mahanta, Jasvinder Kaur, Krishnendu Mondal, Nabeel Ahmad, Inderjeet Singh, Gaurav Sharma, Vipin Gupta

## Abstract

Monkeypox is a viral zoonosis with symptoms that are reminiscent to those experienced in previous smallpox cases. GSAID databases (Global Initiative on Sharing Avian Influenza Data) was used to assess 630 genomes of MPXV. Six primary clades were inferred from the phylogenetic study, coupled with a lesser percentage in radiating clades. Individual clades that make up various nationalities may have formed as a result of a particular SNP hotspot type, which may have mutated in a particular population type.

The most significant mutation, based on a mutational hotspot analysis, was found at G3729A and G5143A. The gene ORF138, which encodes the Ankyrin repeat (ANK) protein, was found to have the most mutations. This protein is known to mediate molecular recognition via protein-protein interactions. It was shown that 243 host proteins interacted with 10 monkeypox proteins identified as the hub proteins E3, SPI2, C5, K7, E8, G6, N2, B14, CRMB, and A41 through 262 direct connections. The interaction with chemokine system-related proteins provides further evidence that the human protein is being suppressed by the monkey pox virus in order to facilitate its survival against innate immunity.

A few FDA-approved molecules were likely used as possible inhibitors after being researched for blocking F13, a significant envelope protein on the membrane of extracellular versions of virus. A total of 2500 putative ligands were docked individually with the F13 protein. The F13 protein and these molecules’ interaction may help prevent the monkey pox virus from spreading. As a result, after being confirmed by experiments, these putative inhibitors might have an impact on the activity of these proteins and be utilised in monkeypox treatments.

## Introduction

Monkeypox has emerged as a new global threat during the COVID-19 pandemic period. The virus that causes this rare zoonotic disease was restricted to just affecting animals when it was first identified in a Danish laboratory in 1958 in monkeys (Brown and Leggat, 2016). It slowly crossed species boundaries, and the first human instance was reported in 1970 in a young child in the Democratic Republic of the Congo (www.who.int) (WHO report, 1980; Ladnyi and Breman, 1978). The illness is caused by the Monkeypox virus (MPXV), a member of genus *Orthopoxvirus* in the Poxviridae family and subfamily Chordopoxvirinae (Pauli et al., 2010). The disease presents clinically in a way that is similar to smallpox. The fact that human monkeypox has returned years after smallpox, a similar orthopoxvirus illness eliminated in 1980, raises concern. After 40 years of no case being documented, monkeypox may have returned owing to several reasons, including an increasing unvaccinated population against smallpox, greater exposure to and interactions with forest animals brought on by deforestation, and population migration (Simpson et al., 2020).

This virus takes 3 to 17 days to incubate, during which time the infected person shows no symptoms (Thornhill et al., 2022; www.cdc.gov.in). The signs often appear three weeks after exposure and include rashes, primarily on and around the genitalia and anus, and may also affect other parts of the body. The rashes might itch and resemble pimples or blisters. Fever, headache, chills, enlarged lymph nodes, sore throat, fatigue, and others could also be present. The disease, however, has a 3-6% fatality rate (www.who.int) and is less contagious and less severe than smallpox. The disease was predominantly confined to regions of central and west Africa with tropical rainforests. However, the lack of a worldwide “one health strategy for disease prevention and treatment” has prevented the disease from being controlled in endemic regions of Africa. Due to the disease’s spread to 110 countries with more than 80 thousandcases that have been reported globally as of December 18, 2022 (https://www.cdc.gov/poxvirus/monkeypox/response/2022/world-map.html), it has become an issue of global public health concern. Very little is currently known about the epidemiology and present trajectory of the virus. Thus, international sequencing attempts to describe the outbreak causing MPXV to pinpoint its lineage and trace its spread started right away. Genome information will also provide data on the genetic diversity and phenotypic characteristics of the virus, which will be relevant for steering research, prophylaxis, and diagnostics.

Although there are several studies that have appeared ever since the emergence of Monkey Pox Virus and it has been reflected that the mutations have been recorded at both the genomic and protein level that drives the evolutionary changes, demanding a detailed study of Monkey Pox mutations to understand its successful invasion and infection. To unveil this, we rendered 630 genomic sequences, deciphering their phylogenetic relationships and tracing their SNPs at nucleotide levels. We further extended the study to understand the mechanism of host immunity evasion by host-pathogen interaction (HPI) by confirming their interactions with host proteins to understand the functional profile of host proteins directly affected by the viral proteins. Further it has been reportedly found that tecovirimat has been the only FDA approved anti-viral drug found effective against MPOX infection. In this study, we have screened ∼2500 FDA approved molecules (on request from DrugBank and not available in public databases) using *in-silico* molecular docking that can be formulated and purposed for monkey pox treatment.

## Material and Methods

### Selection of genomes, annotations, and phylogeny construction

The selected MPXV genomes were retrieved upon request from GSAID public database. As of July 2022, only 629 MPXV genomes were available in the database (see Data Set S1, sheet 1, in the supplemental material) that were incorporated in this study for comparative analyses. Completely annotated reference genome with accession no NC_063383 was selected from NCBI virus genome database. Finally, a total of 630 complete genome sequences were selected for this study to determine MPXV phylogeny and comparative genome analyses.

### Genotyping based on SNP

To detect nucleotide variations among 629 genomes of MPXV, multiple sequence alignment was performed against the reference genome. The nucleotide changes were calculated as point variations and recorded. The interpolation and visualization were plotted using computer programs in Python.

### Data and computer programs

The genomic analytics is performed using Python and Biopython libraries (83). The computer programs and the updated SNP profiles of MPXV isolates are available upon request.

### Host-Pathogen interactions and functional annotations

To find the host-pathogen interaction (HPI), we subjected MPXV protein sequences to host-pathogen interaction databases such as UniProt (ref), Viruses STRING v10.5 (Cook et al., 2018) and HPIDB3.0 (Ammari et al., 2016) to predict their direct interaction with humans as the principal host. In these databases, the virus-host interaction was imported from different PPI databases like MintAct (Orchard et al., 2014), IntAct (Orchard et al., 2014), HPIDB (Ammari et al., 2016), and VirusMentha (Calderone et al., 2015). It searches protein sequences using BLASTP to retrieve homologous host/pathogen sequences. For high-throughput analysis, it searches multiple protein sequences at a time using BLASTp and obtains results in tabular and sequence alignment formats (Kumar and Nanduri, 2010). The HPI network was constructed and visualized using Cytoscape v3.9.1 (Shannon et al., 2003). The MPXV hub proteins from the host-pathogen interactome were identified using the Maximal Clique Centrality (MCC) algorithm in the CytoHubba plugin (Chin et al., 2014) in Cytoscape v3.9.1. Further, the human proteins interacting with individual viral proteins were subjected to functional annotation. For this, Gene ontology (GO) analysis was performed using ClueGo (Bindea et al., 2009), selecting the Kyoto Encyclopedia of Genes and Genomes (KEGG) (Kanehisa et al., 2016), Gene Ontology biological function database, and Reactome Pathways (Fabregat et al., 2018) databases. The ClueGo parameters were as follows: Go

Term Fusion selected; pathways or terms of the associated genes, ranked based on the P value corrected with Bonferroni stepdown (P values of <0.05); GO tree interval, all levels; GO term minimum number of genes, 3; threshold, 4% of genes per pathway; kappa score, 0.42. Gene ontology terms are presented as nodes and clustered together based on the similarity of genes corresponding to each term or pathway.

### Structure Modelling and validation

The 3D structure of F13 viral protein was predicted using *de-novo* modelling algorithm of I-Tasser server (Yang et al., 2015). The models were validated based on the quality score and the Ramachandran plot was predicted using PDBsum. The 3D structure was further subjected to structure validation using ProSAweb (Wiederstein et al., 2007) Protein Quality Predictor (ProQ) (Cristobal et al., 2001) and Research Collaboratory for Structural Bioinformatics (RCSB) (Burley et al., 2019) validation server and Z-score and LG score were calculated.

### Ligand Preparation

The FDA-approved molecules (n=2500) present in the drug bank (Law et al., 2014) were used for this study. The chemical structures of these drugs were retrieved in SDF format from the DrugBank (https://go.drugbank.com/) (on request). The structures were converted into mol2 format by Avagardo and Open Babel software (O’Boyle et al., 2011). The resultant files were then used for carrying out molecular docking. Additionally, The FDA-approved molecules (n=2500) provided by were used for this study.

### Molecular Docking and Simulations

The multi-docking for 2500 ligands and 6 NAPs were carried out using PyRx software (https://pyrx.sourceforge.io/). Protein files were prepared using Biovia Discovery Studio by deleting water molecules and addition of polar hydrogens (Tanwar and Purohit, 2019). Energy was also checked for each structure. AutoDock Vina, a plugin of PyRx was used for molecular docking. PBD file of the proteins molecules were converted into macromolecule (pdbqt file). The ligand molecules (sdf or mol2 format) were prepared using the OpenBabel, plugin software of PyRx by energy minimizing (Rosário-Ferreira et al., 2021). Before docking, the molecules were prepared and converted for autodock. Blind docking was performed between prepared macromolecules and ligand molecules using AutoDock4 Vina Wizard.

## Results and Discussion

### Phylogenetic relationship between different MPXV strains and their spread

The availability of genomic sequences in public databases and highlighted surveillances across the globe the assessment of mutations in monkeypox were calculated based on the single nucleotide polymorphism (SNPs). As mentioned in the methods 630 (629+reference) genomes of Monkeypox were assessed through GSAID databases. The phylogenetic analysis deduced six major clades along with a smaller fraction in radiating clades. The formation of individual clades constituting different countries may be attributed to the type of specific SNP hotspot type which might have occurred due to the mutation in specific population type. Moreover, it has been previously reported that travel history is one of the main reasons for abrupt transmission and wide spread among different geographical locations for viral infections (Gupta et al. 2021, Kumar et al., 2021). However, such analysis with the help of artificial intelligence algorithm may be used to track down the spread of major SNPs favored in specific geographical locations. The independent clades radiating out from main clade highlighted the sporadically present SNPs in different genomic locations signifying that not all mutations occurred are uniformly favored in MPXV genome. These radiating locations can also be representing accessory mutations which might be selected by nature for survival of MPXV in varied environmental and host condition.

### Mutational Hotspots

Multiple sequence alignment was performed for all 630 MPXV genomes using the reference strain (NC_063383). We identified some sites with >90% mutation rates as compared to the reference. It was seen that 67 sites harboured >15% mutation probability. 851 sites had 2-15% mutation probability and <2% mutation probability was found in 4,550 sites. The most significant mutation was identified at G3729A and G5143A. ORF138 which codes for Ankyrin repeat (ANK) protein known to mediate molecular recognition via protein-protein interactions, was found to have maximum mutations. Highest mutational frequency was observed (transitions and transversions) from A to G (185) and T to C (175) followed by G to A (153) and C to T (152). Maximum AG mutations were found in genes lying between location 180000 to 190000 that harbour proteins including ORF179, ORF264, ORF180, and ORF89. ORF264 is a membrane glycoprotein whereas the function remains unknown for other proteins (Figure 2).

**Figure 1:**
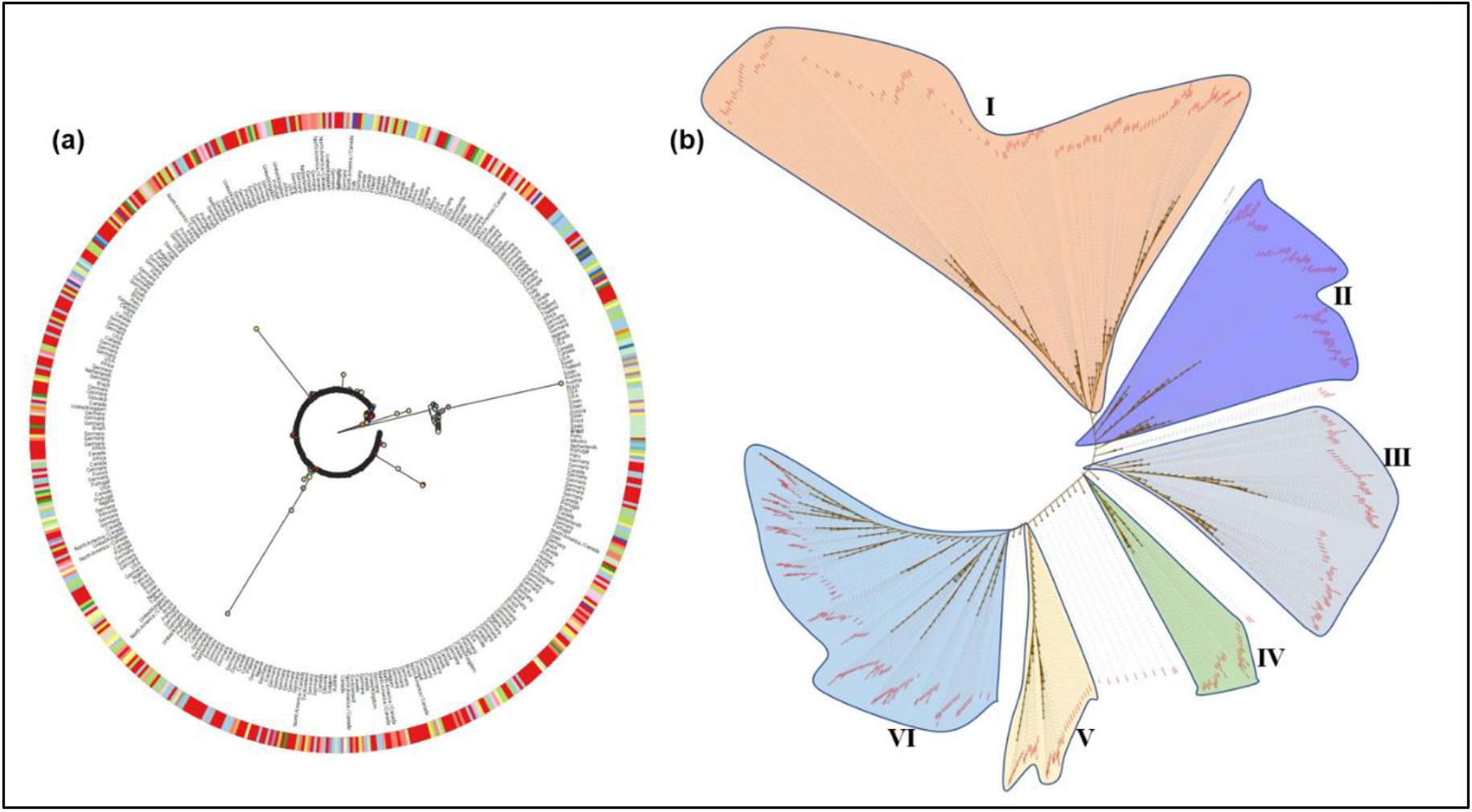
Phylogenetic network of 630 MPXV genomes. (a) Nucleotide-based phylogenetic analysis of MPXV isolates using the maximum likelihood method based on the Tamura-Nei model. (b) The map is diverged into six major clades (clades I to VI) representing variation in the genomes. The coloured circle represents the country of origin of each isolate.

**Figure 2:**
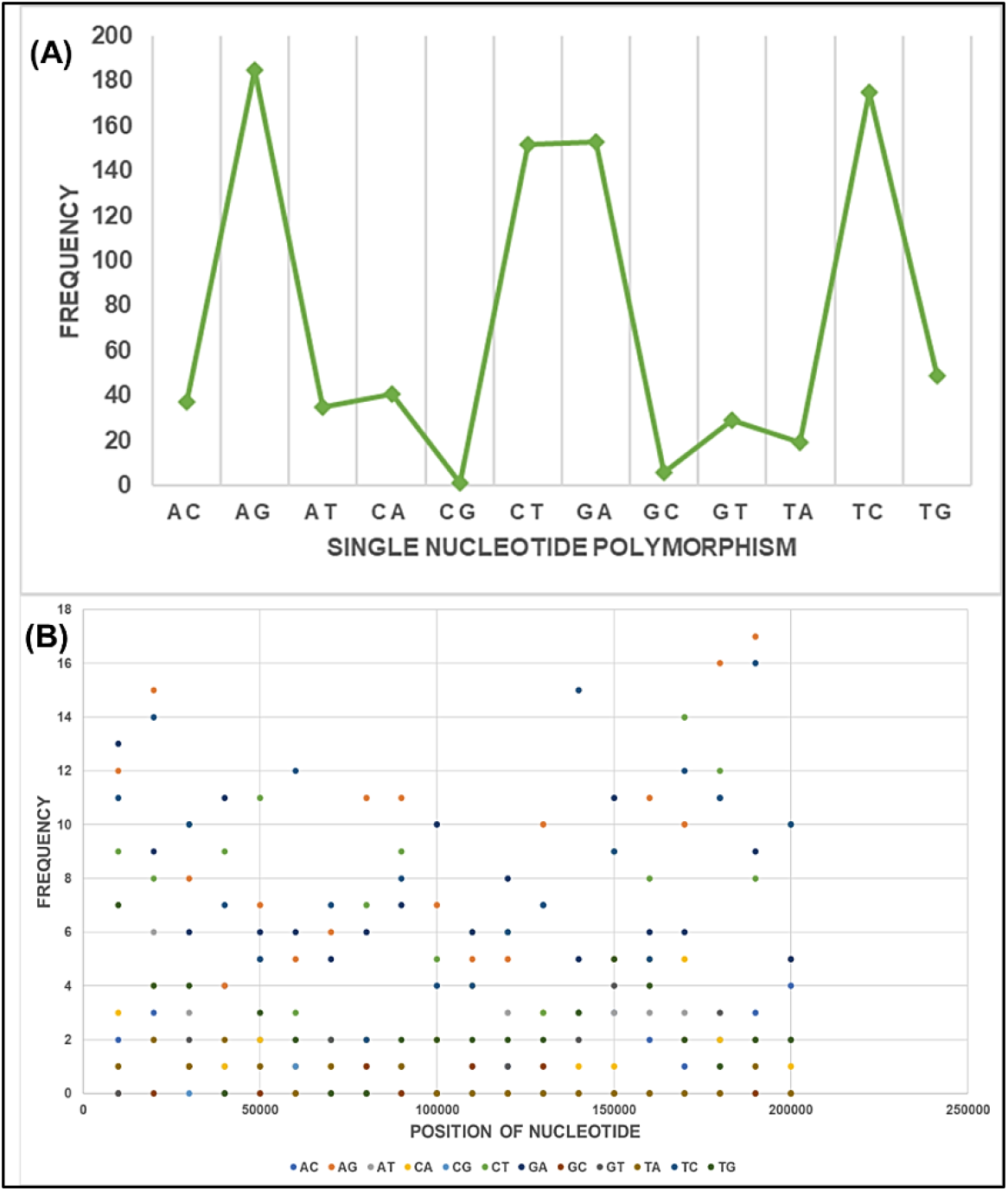
Distribution of SNP mutations of MPXV isolates from the globe. (A) Frequency-based plot of 12 possible SNP mutations across 630 genomes. (B) Frequencies of the single SNP mutations with locations on the genome. The nucleotide positions are based on the reference genome of MPXV.

### Monkeypox-Human Interactome

The interactions between virus and host proteins are key to maintaining viral life cycle within the host and the study of these interactions promises to elucidate host mechanisms that viruses evade in order to replicate. In this study we performed a protein-protein interaction study using the complete proteomes of Monkeypox virus and human to determine the key virus proteins that interact with those of humans upon infection. The interactome contained 385 total proteins, of which 342 were human proteins and 43 were of virus origin (Figure 3A). We identified 10 Monkeypox proteins as the hub proteins using Maximal Clique Centrality (MCC) algorithm in CytoHubba (Chin et al., 2014) within Cytoscape. These hub proteins - E3, SPI2, C5, K7, E8, G6, N2, B14, CRMB and A41 interact with 243 host proteins (nodes) through 262 direct interactions (edges) (Figure 3A). Limited by their small genomes, viruses are known to interact with and regulate several host cellular pathways through a small number of proteins.

**Figure 3:**
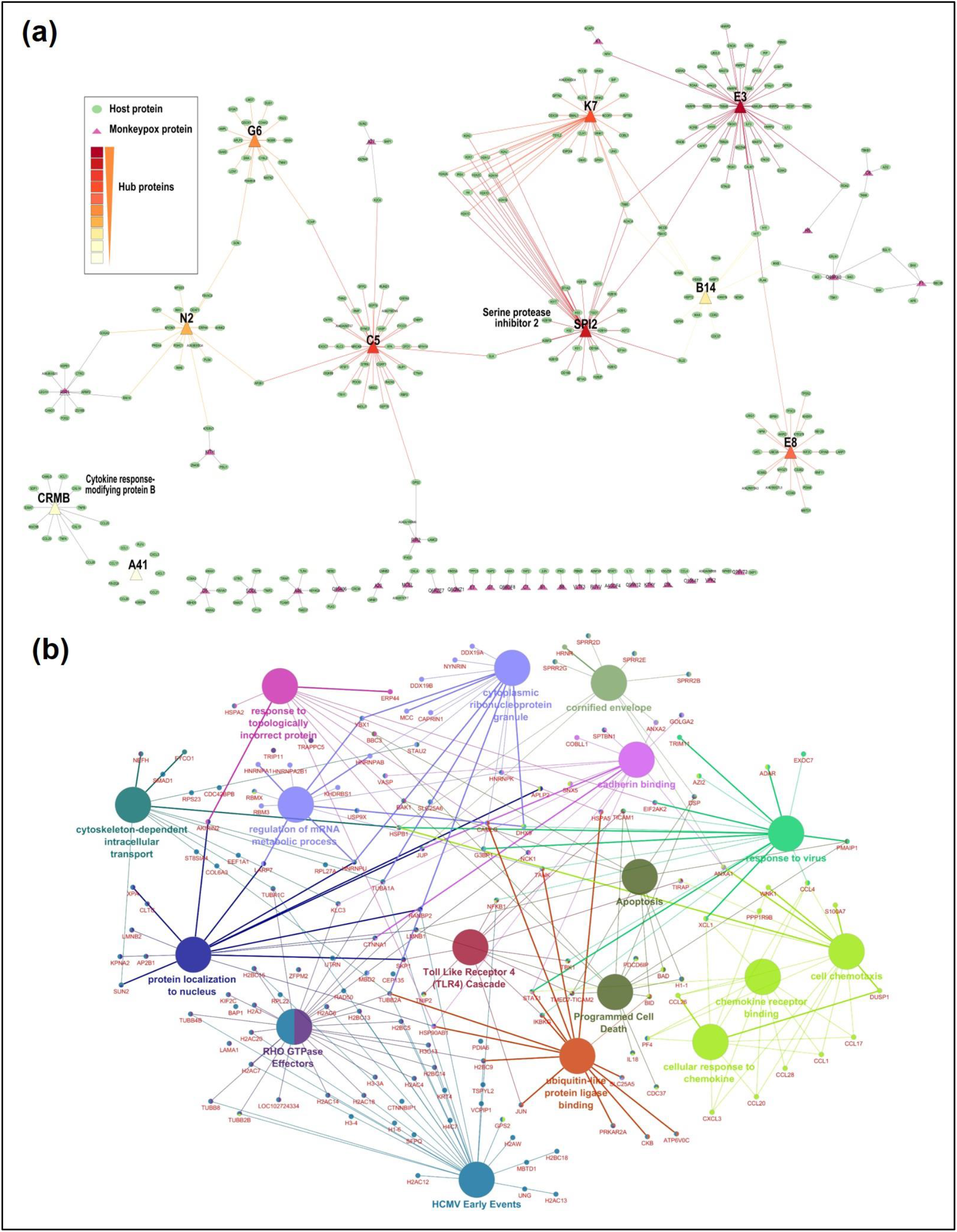
(A) Host-pathogen interaction of MonkeyPox viral proteins and human proteins. Nodes represent proteins, while lines/edges represent interaction. Triangles (red) represent viral proteins found to be directly interacting with the human proteins (green). The hubs (E3, SP12, C5, K7) were found interacting with maximum viral proteins. (B) Gene ontology (GO) analysis was performed for host proteins using the ClueGo Cytoscape app against database KEGG, the Gene Ontology—biological function database, and Reactome pathways. ClueGo parameters were set as follows: Go Term Fusion selected; P values of #0.05; GO tree interval, all levels; kappa score of 0.42.

The E3 hub protein was found to interact with 50 human proteins, SPI2 interacted with 38 host proteins, C5 with 37, K7 with 35, E8 with 24, G6 with 19, N2 and B14 with 18, crmb with 12 and A41 with 11 human proteins (Figure 3A). The E3 protein of vaccinia virus is shown to play critical role in evading host immune responses by inhibiting antiviral interferon (IFN) signaling (Arndt et al., 2015). The SPI2 protein is shown to have antiapoptotic activity (Dobbelstein and Shenk, 1996) and is also involved in inhibiting IFN-B induction along with cytokine response modifier A (CrmA) (Qin et al., 2017). Another MPXV protein C5, identified as the hub forms a component of the monkeypox inhibitor of complement enzymes (MOPICE) and thus acts in immunomodulatory strategies of the virus (Weaver and Isaacs, 2008). The K7 protein identified as the hub protein in MPXV antagonises the host innate immune response by binding Toll-like receptor-adaptor proteins and the DEAD-box RNA helicase DDX3. DDX3 is a key component in the signaling cascade that leads to induction of genes encoding Type I interferons. K7 binding to DDX3 is thought to simply interfere with the downstream signaling and inhibiting NF-κB and interferon regulatory factor 3 activation (Teferi et al, 2017, Benfield et al., 2013). These interactions of MPXV K7 with host DDX3 was confirmed in our interactome study. Another important interaction was between K7 with the SPIR1 protein that is activated within host as a restriction factor upon infection by diverse DNA and RNA viruses (Torres et al., 2022). K7 is also known to inhibit several antiviral proteins by promoting cellular histone methylation during infection as was evident from its interaction with human histone proteins in the interactome (Teferi et al., 2017). K7 also interacted with members of the WNK (with no lysine) family – WNK1, WNK2 and WNK3. The WNK family proteins WNK1 and WNK3 are responsible for interleukin-1 cytokine activation and therefore form vulnerable nodes in the host cellular defence system (Pichlmair et al., 2012). Therefore, their suppression by K7 viral protein is one of the crucial host immune evasion strategies of the monkeypox virus.

Poxviruses encode several viral soluble TNF receptors that inhibit the proinflammatory cytokine via C-terminal domain. These TNF-binding proteins, for example CrmB, that was identified as a hub in the MPXV-human interactome, are known to contribute to the pathogenicity of the poxviruses and have been elucidated in the variola (VARV), monkeypox (MPXV), and cowpox (CPXV) viruses (Shchelkunov et al, 1993; Upton et al., 1991). The other proteins such as E8 plays regulatory roles in viral replication (Zobel et al., 2003); N2, a nuclear localizing MPXV protein is a virulence factor that inhibits the activation of interferon regulatory factor 3 (IRF-3) (Ferguson et al., 2013) and A41 protein binds with the chemokines and affects host immune responses (Bahar et al., 2014). B14 is another poxvirus virulence factor that inhibits the inhibitory κB (IκB) kinase (IKK) complex by preventing ubiquitin-mediated degradation of NFκB inhibitor IkBα (Mohamed and McFadden, 2009) (Figure 3b).

Functional profile of human protein found interacting with the MPOX virus protein include immunity related functions like TLR4 cascade, cytoskeleton dependent intracellular transport protein, cytoplasmic ribonucleoprotein granules, cadherin binding, response to virus, cellular response to chemokine, response to topologically incorrect protein, cornified envelop. Toll-Like Receptor 4 (TLR4) signal pathway plays an important role in initiating the innate immune response (Kuzmich et al., 2017). Cellular cytoskeleton on the other hand provides the basis for intracellular movements such as those that transport the pathogen to and from the cell surface (Bearer et al., 2002). RNA granules are found in all types of eukaryotic cells and tissues and are involved in many aspects of gene regulation, homeostasis and cytopathology. They have been known for their role to restrict poxvirus replication through the formation of antiviral granules (Rozelle et al., 2014). The profile of chemokine expression contributes to shaping the immune response during viral infection, whereas viral subversion of the chemokine system allows the virus to evade antiviral activities of the host (Melchjorsen et al., 2003). DNA viruses like herpesviruses and poxviruses are known to encode proteins that mimic chemokines and chemokine receptors, as a part of strategic evasion to human innate immunity. They encode homologs of chemokine ligands and chemokine receptors, and secreted chemokine binding proteins (CKBPs) to curb chemokine activity (Alcami, 2003). The interaction with proteins linked to chemokine system confirm the fact that monkey pox viral proteins are interacting and suppressing the human protein to make its survival easy against the innate immunity (Alcami and Lira, 2010).

### Repurposing and molecular docking of drugs and target proteins

The 3-Dimensional structure of F13 protein was predicted using *de-novo* modelling using I-Tasser. The selected structure had a quality score in the reported range and are represented in Figure 4. Ramachandran score was calculated and was in range with more than 90% of residue in the favoured. The protein’s active site is an essential criterion while performing molecular docking. The active site was predicted using ScanProtSite. The sites were taken into consideration while performing molecular docking. In order to find out the potential inhibitors, the *in-silico* molecular docking studies were carried out through AutoDock Vina. The F13 protein was individually docked with the mentioned 2500 ligand. Binding affinities were calculated for each docking (Supplementary Table 1). The binding affinity less than -10 and rnsd/ub and rmsd/lb values 0 was selected for analysis. Out of more than 2500 docking results, 7 docking between F13 and 7 ligands (Fig 4; Table 3) were found to have binding affinity less than -10 making it strong interaction. Till date only tecovirimat is a known and potential drug that target monkey pox virus. F13 is the major envelope protein on the membrane of extracellular forms of virus. This protein has been reported from the vaccinia virus, encoded by the F13L gene, and is known to be conserved across the subfamily Chordopoxvirinae. Moreover, it is critical among genus Orthopoxvirus members to produce the wrapped form of virus that is required for cell-to-cell spread. The interaction between the molecule and the F13 protein will promote blocking of the spread of monkey pox virus. Therefore, these potential inhibitors might affect these proteins activity and can be potentially used in monkeypox therapeutics after confirmation with experimentations.

**Table 3.** Binding energy of molecules and F13 protein from monkey pox virus.

| Ligand | Ligand ID | Binding energy | rsmd |
| --- | --- | --- | --- |
| structures_uff_E=23331.24_out | #1 | -13.1 | 0 |
| structures_uff_E=14383.50_out | #2 | -12 | 0 |
| structures_uff_E=9521.68_out | #3 | -12 | 0 |
| structures_uff_E=17011.61_out | #4 | -11.8 | 0 |
| structures_uff_E=7659.67_out | #5 | -11.8 | 0 |
| 320 | #6 | -11.3 | 0 |
| structures_uff_E=11743.10_out | #7 | -11.1 | 0 |

**Figure 4:**
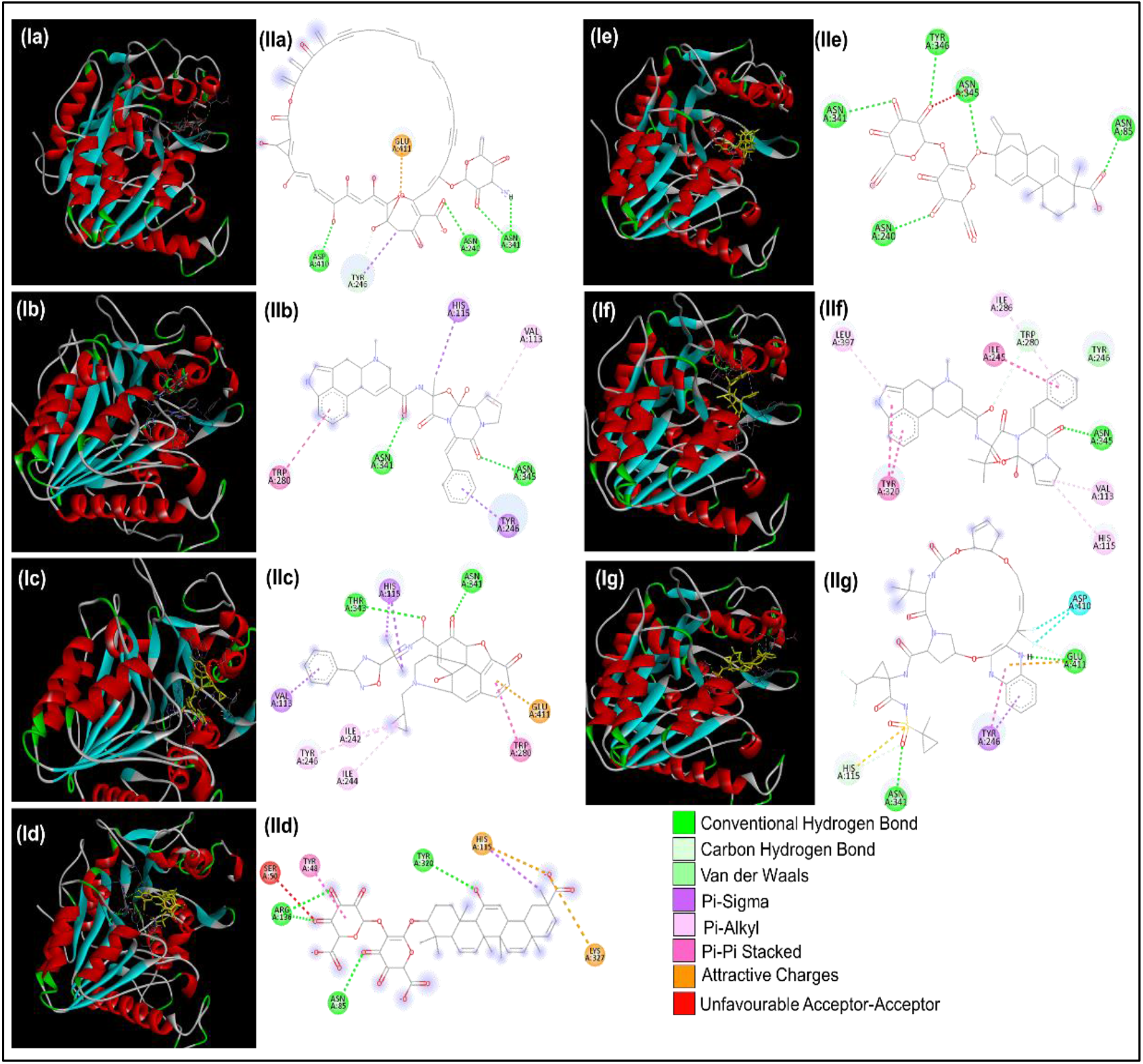
Molecular docking of MPXV protein F13 with FDA approved molecules namely Ligand 1-7 (as named in table 3). The docked conformation of the Ligands depicting the possible and detailed interactions with various amino acids of NAPs.

## Conclusion

This genomic and proteomic survey of Monkey Pox Viral strains using subsets of populations isolated from different countries investigate their mutation prevalence and host-pathogen interactions. The viral phylogenetic network with six clades (I to VI) provided a landscape of the current stage of epidemic where non-directional approach is being followed. In this study we also performed a protein-protein interaction study using the complete proteomes of Monkeypox virus and human to determine the key virus proteins that might interact with those of humans upon infection. Ten hub proteins - E3, SPI2, C5, K7, E8, G6, N2, B14, CRMB and A41 were identified to interact with 243 host proteins. Majority of the interacting host partner proteins were found to be related to immunity related functions confirming the attack of virus directly to immunity of the host. Till date only tecovirimat is a known and potential drug that target monkey pox virus. FDA approved molecules were investigated to repurpose for blocking F13, a major envelope protein on the membrane of extracellular forms of virus revealing the probable use of a few molecules as potential inhibitors. This multiomics approach using genomics, proteomics, interactomes, and systems and structural biology has provided us an opportunity for better understanding of monkey pox strains and its variants.

## Data availability

Complete set of sequences for full-length genomes and proteomes of MPXV virus used in the study are available at GSAID database (https://gisaid.org/hmpxv-phylogeny/).

